# Analysis and subsequent molecular docking of selected phytochemicals with SLC6A3 and SLC6A4 as potential therapeutic agents for Obsessive-Compulsive Disorder (OCD)

**DOI:** 10.1101/776922

**Authors:** Swarn S. Warshaneyan, Prachi Srivastava

## Abstract

Obsessive-Compulsive Disorder (OCD) is one of the most problematic disorders of the brain. It lies in the category of moodomics disorders and can be identified by the presence of recurring intrusive thoughts (“compulsions”) and urges to do certain things repeatedly (“obsessions”), with a definite lack of control on these tendencies and their intensity affecting the patient’s daily life. It is known to be associated with not only anxiety disorder but also depression, with its cause being unknown, although possessing genetic components. Several separate methods are being attempted to develop an effective therapy for OCD but this field of research requires more focus. In the present course of research, there is an attempt to signify the molecular analysis approach by testing potential drugs computationally with previously identified targets, i. e. dopamine active transporter (DAT or SLC6A3) and sodium-dependent serotonin transporter (SERT or 5-HTT or SLC6A4) – two proteins which have been observed playing a major role in the manifestation of OCD symptoms. Their 3D structure prediction has already been done in the previous course of study. In continuation to the previous study, there is an attempt to screen out the potentially useful phytochemicals against this moodomics disorder. Several potentially useful phytochemicals are known to associate with SLC6A3 and SLC6A4, which have been identified through the text mining of various publications. First of all, these phytochemicals are analysed through the use of Molinspiration software followed by selection of the ones suitable as drug candidates using Lipinski’s Rule of five and finally, best docking configurations identified using AutoDock 4 in order to obtain potential therapeutics for OCD patients. Nuciferine, Epicatechin (EC), Retinal, Aphidicolin and Salvinorin A are showing good docking scores but among all these phytochemicals, Nuciferin, an extract of Indian Lotus plant, appears to be the one most suitable compound on the basis of binding energy calculations. Hence, we can say this could be a good revealing therapeutic agent for this disorder.

## INTRODUCTION

Plaguing about 2.3% of the human population for at least some time in their lives, Obsessive-Compulsive Disorder (OCD) is probably one of the most problematic disorders of the brain, sharing common points and associated with not only anxiety disorder but also depression, tic disorder and increased suicide risk [1][2][3]. It is a moodomics, i. e. mood and emotion omics, disorder and can be identified by the presence of symptoms which include recurring intrusive thoughts (known as “compulsions”) and urges to do certain things repeatedly (known as “obsessions”), with a definite lack of control on these tendencies and their intensity affecting the patient’s daily life, in a clearly negative manner [4]. The cause is unknown but possesses genetic components, identifiable by the fact that both identical twins are affected more often as compared to both non-identical twins [2][4]. Although multiple different methods are being attempted in order to develop an effective therapy for OCD, still this field of research requires more focus and deeper investigation. In the present course of research, there is an attempt to signify the molecular analysis approach through bioinformatics methods by testing potential drug candidates computationally with previously identified targets, i. e. dopamine active transporter (DAT or SLC6A3) and sodium-dependent serotonin transporter (SERT or 5-HTT or SLC6A4) – two proteins which have been observed playing major roles in the manifestation of OCD symptoms through their association with two major neurotransmitters – dopamine and serotonin [5][6] [7][8]. Their 3D structure prediction has already been performed in the previous course of study. In continuation to the previous study, there is an attempt to screen out the potentially useful phytochemicals against this moodomics disorder. Several potentially useful phytochemicals are known to associate with SLC6A3 and SLC6A4, which have been identified through the text mining of various publications documenting the association between known phytochemicals, dopamine transporter and serotonin transporter [9][10][11][12][13][14][15]. As the first step, a list of phytochemicals is prepared by going through various papers and then their canonical SMILES line notations are retrieved from PubChem Compund, which are used for calculating the molecular properties of these phytochemicals through the use of Molinspiration, followed by selection of the ones suitable as drug candidates using Lipinski’s Rule of five and finally, the ones which are selected have their best docking configurations identified using AutoDock v4.2.6 in order to obtain potential therapeutics for OCD patients [16][17]. In this step, a total of 9 phytochemicals were used for molecular docking studies with both dopamine transporter and serotonin transporter. The ones which produced the best scoring results were found out to be Nuciferine, Epicatechin (EC), Retinal, Aphidicolin and Salvinorin A, which are showing good molecular docking scores for both dopamine transporter and serotonin transporter; while the top three results were that of Nuciferine, Retinal and Salvinorin A.

## MATERIALS & METHODS

This work involved the utilization of previously produced 3D structures (homology models) of the target proteins, i. e. dopamine transporter and serotonin transporter (through the use of Swiss-Model workspace, which were validated using Verify3D from SAVE Server v5.0), for the purpose of molecular docking with phytochemicals (selected using Lipinski’s Rule of five) from a list made by text mining relevent publications. This step was performed using AutoDock v4.2.6 in order to obtain the best configuration for each of the 9 phytochemical ligands from the previous step (out of 10 possible configurations for each phytochemical ligand).

### Ligand Selection

PubMed (https://www.ncbi.nlm.nih.gov/pubmed) was used for retrieving papers which showed an association between certain phytochemicals with dopamine transporter and serotonin transporter, which in turn allowed a list of around 20 to 30 phytochemicals to be made but a better look into the list resulted in the exclusion of several toxic phytochemicals and eventually, 14 non-toxic phytochemicals were selected as ligands through text mining of various publications. The number was further shaved down to 9 ligands through the application of Lipinski’s Rule of five, with the final ligands being as follows: Alpha-Pinene, Aphidicolin, Epicatechin (EC), Forskolin, Geranylgeranyl pyrophosphate, Nuciferine, Phytol, Retinal and Salvinorin A.

### Ligand Retrieval

The ligand structures and their canonical SMILES line notations were retrieved from the PubChem Compound (https://www.ncbi.nlm.nih.gov/pccompound). The ligand structures were retrieved as SDF (Structure Data Format) files and were converted to PDBQT (Protein Data Bank, Partial Charge (Q) & Atom Type (T)) format through the use of open-source chemistry software tool Open Babel v2.4.1 for the purpose of being used with AutoDock v4.2.6, while their molecular properties were calculated through Molinspiration by using their canonical SMILES line notations as inputs for the program [18].

### Molecular Docking Analysis

AutoDock v4.2.6, a suite comprising of automated molecular docking tools, was utilised for the molecular docking analysis of the selected ligands with dopamine transporter and serotonin transporter. The PDBQT files of both proteins were also generated through this software (using their previously created PDB files as inputs). Once the molecular dockings were finished and 10 configurations for each phytochemical-ligand complex were generated for all the ligands using the software’s in-built Lamarckian Genetic Algorithm feature, text files of scoring results were also produced for the purpose of manual comparative analysis. The final phytochemical-ligand complex analysis was performed with the help of Discovery Studio 2019 [19].

## RESULTS & DISCUSSION

### Structure Prediction and characterization of target proteins SLC6A3 and SLC6A4

Since at the time of the preceding as well the current study, the 3D structures of target proteins, i. e. dopamine active transporter (DAT or SLC6A3)(shown in Figure 2a) and sodium-dependent serotonin transporter (SERT or 5-HTT or SLC6A4)(shown in Figure 2b), weren’t available on PDB, they were generated through homology modeling using Swiss-Model workspace, while their structural details such as the percentage of alpha helices, extended strands, beta turns, random coils etc. alongwith their molecular properties, such as aliphatic index, instability index, GRAVY, estimated half-life, extinction coefficients, number of charged residues, atomic composition etc. were analysed using SOPMA (depicted in Table 1a) and ProtParam (depicted in Table 1b and Table 1c) respectively, through their sequences which were obtained in FASTA format from UniProt [20][21][22]. By determining these values, we arrived at the conclusion that both the target proteins are quite similar in structure and function. Both the 3D structures were then validated with the help of Verify3D at SAVE Server v5.0 and each received a passing score [23]. In order to receive a “Pass” from Verify3D, a model needs to get at least 80% of its residues to receive an averaged 3D-1D score greater than 0.2 and for the SLC6A3 model, the result was 82.14%, while for the SLC6A4 model, it was 88.75%. Both the models were then visually analysed using the software tool Chimera v1.13.1 by UCSF [24].

**Figure 1:**
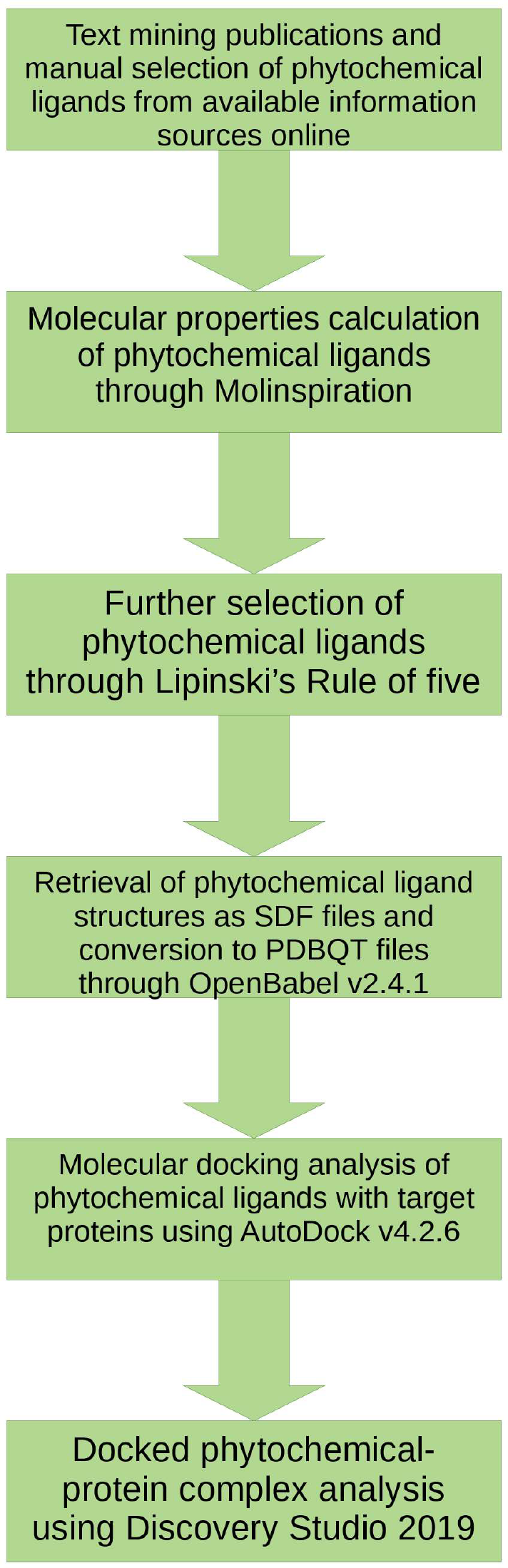
Flowchart depicting the workflow in a schematic manner.

**Figure 2(a):**
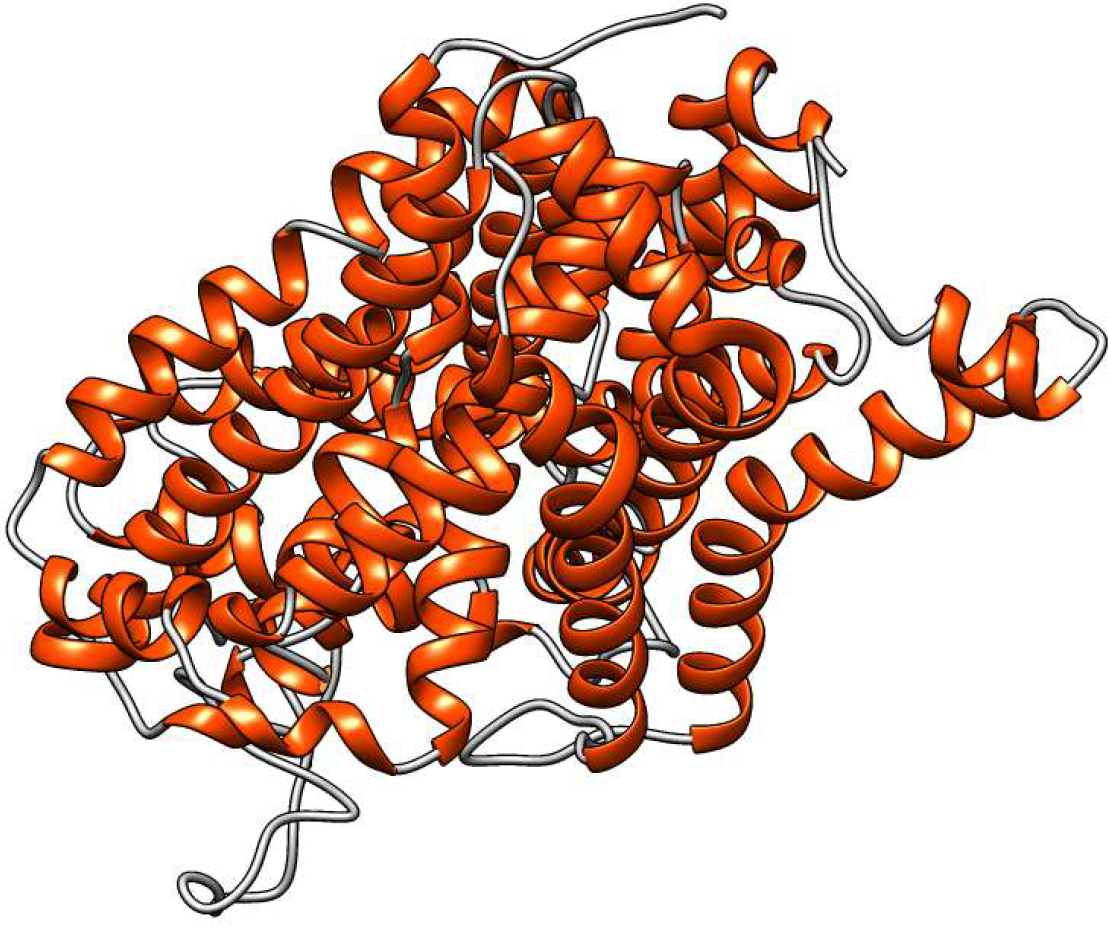
SLC6A3 model visualised using UCSF Chimera (helices: orange, strands: purple, coils: grey)

**Figure 2(b):**
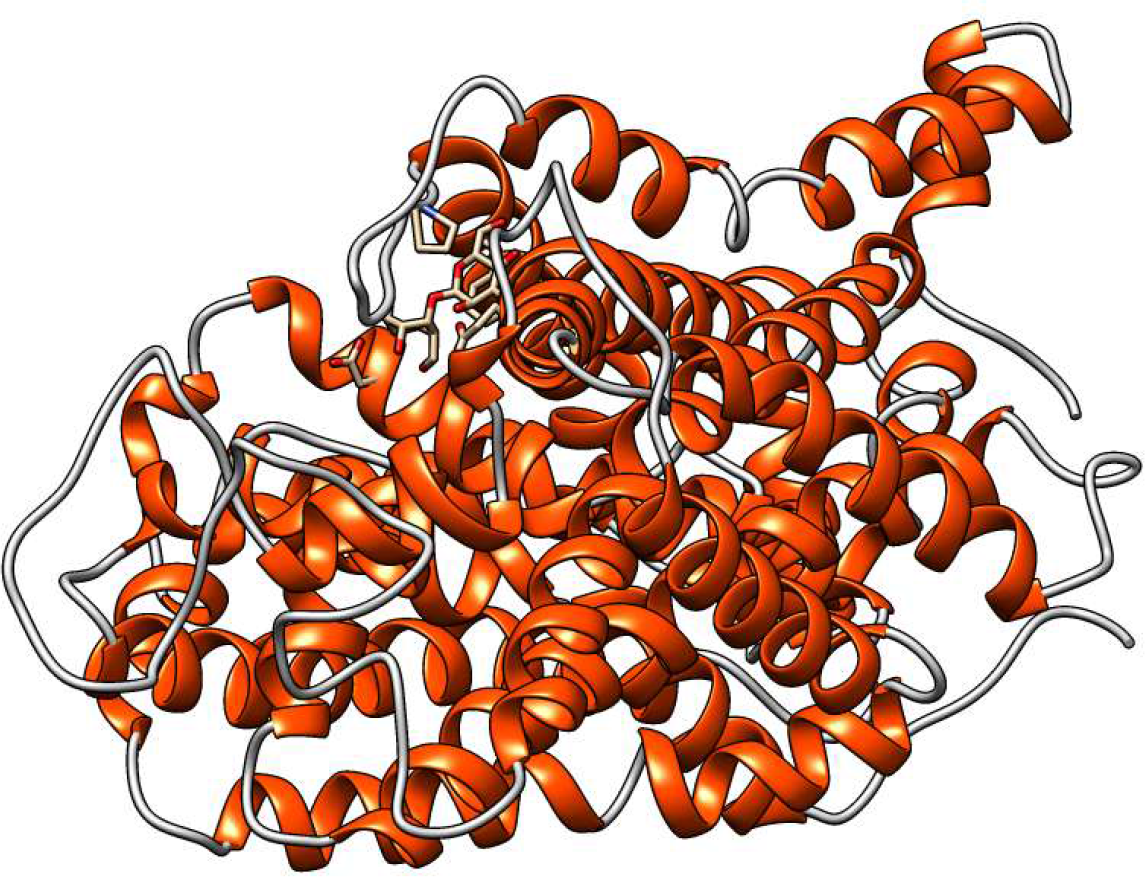
SLC6A4 model visualised using UCSF Chimera (helices: orange, strands: purple, coils: grey)

**Table 1(a):** Structural details of SLC6A3 and SLC6A4 computed through SOPMA.

| Serial Number | Target Protein | Sequence length | Alpha helix (Hh) | Extended strand (Ee) | Beta turn (Tt) | Random coil (Cc) |
| --- | --- | --- | --- | --- | --- | --- |
| 1. | SLC6A3 | 620 | 241 is<br>38.87% | 119 is<br>19.19% | 40 is 6.45% | 220 is<br>35.48% |
| 2. | SLC6A4 | 630 | 232 is<br>36.83% | 116 is<br>18.41% | 48 is 7.62% | 234 is<br>37.14% |

**Table 1(b):** Physiochemical parameters of SLC6A3 and SLC6A4 computed through ProtParam.

| Protein | SLC6A3 | SLC6A4 |
| --- | --- | --- |
| Accession Number | Q01959 | P31645 |
| Primary Sequence Length | 620 | 630 |
| Molecular Mass | 68494.91 | 70324.86 |
| Theoretical pI | 6.46 | 5.89 |
| Negatively Charged Residues | 44 | 45 |
| Positively Charged Residues | 41 | 40 |
| Extinction Coefficient (Cys forms Cystine) | 130010 | 151815 |
| Extinction Coefficient (Cys reduced) | 129260 | 150690 |
| Instability Index (II) | 30.85 | 33.13 |
| Aliphatic Index (AI) | 105.32 | 100.44 |
| Grand Average of Hydropathicity | 0.499 | 0 |

**Table 1(c):**
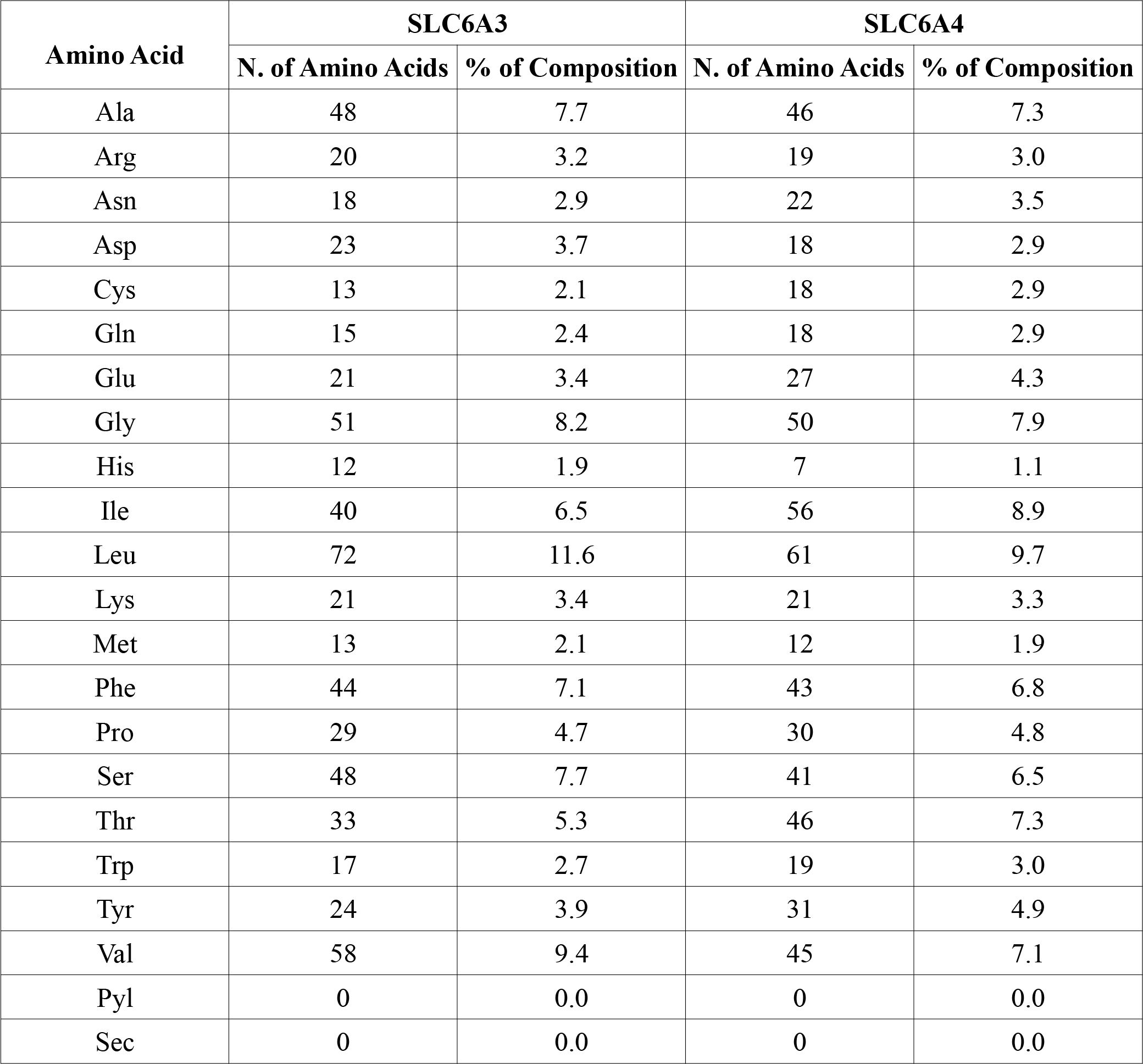
Amino Acid compositions of SLC6A3 and SLC6A4 computed through ProtParam.

### Construction of Virtual Library of phytochemicals

A number of potential phytochemical ligands were identified by performing text mining of relevant research papers noting the association between certain phytochemicals and our target proteins, namely, dopamine-transporter and serotonin-transporter. After looking into the information already available about these phytochemicals (around 20 to 30 in number), all those which have been observed to have known toxic effects on the human body, such as MPTP (a precursor to the neurotoxic MPP+), were dropped, leaving us with a list made up of 14 phytochemicals. The canonical SMILES line notations of these phytochemicals were retrieved from PubChem Compound database and were used for calculating molecular properties through Molinspiration (depicted in Table 2). The values generated were used with Lipinski’s Rule of Five to further reduce to number of phytochemicals suitable as ligands to 9. Now the phytochemical ligand structures were again obtained from the PubChem Compound database to be used for the purpose of molecular docking and complex analysis with SLC6A3 and SLC6A4 using AutoDock v4.2.6 and Discovery Studio 2019 respectively.

**Table 2:** Molecular properties of phytochemical ligands calculated through Molinspiration, the number of Lipinki’s Rule of Five violations (nviolations) and their respective PubChem CIDs.

| S. N. | Phytochemical | miLogP | TPSA | natoms | Molecular Weight | nOH | nOHNH | nviolations | nrotb | Volume | PubChem CID |
| --- | --- | --- | --- | --- | --- | --- | --- | --- | --- | --- | --- |
| 1. | Alpha-Pinene | 3.54 | 0 | 10 | 136.24 | 0 | 0 | 0 | 0 | 151.81 | 6654 |
| 2. | Aphidicolin | 1.85 | 80.91 | 24 | 338.49 | 4 | 4 | 0 | 2 | 335.80 | 457964 |
| 3. | Epicatechin (EC) | 1.37 | 110.37 | 21 | 290.27 | 6 | 5 | 0 | 1 | 244.14 | 14332898 |
| 4. | Forskolin | 1.71 | 113.29 | 29 | 410.51 | 7 | 3 | 0 | 3 | 387.43 | 47936 |
| 5. | Geranylgeranyl pyrophosphate | 5.22 | 113.29 | 29 | 450.45 | 7 | 3 | 1 | 14 | 421.44 | 447277 |
| 6. | Nuciferine | 3.45 | 21.71 | 22 | 295.38 | 3 | 0 | 0 | 2 | 281.44 | 10146 |
| 7. | Phytol | 66.76 | 20.23 | 21 | 296.54 | 1 | 1 | 1 | 13 | 349.38 | 5280435 |
| 8. | Retinal | 6.38 | 17.07 | 21 | 284.44 | 1 | 0 | 1 | 5 | 307.57 | 638015 |
| 9. | Salvinorin A | 2.8 | 109.13 | 31 | 432.47 | 8 | 0 | 0 | 5 | 386.57 | 128563 |

### Molecular docking studies with SLC6A3 and SLC6A4

The molecular docking were performed with the 9 phytochemical ligands and the 2 proteins using AutoDock v4.2.6 to generate a total of 18 results, with 10 possible configurations in each result. The best configuration was selected for each ligand through the process of elimination by using free binding energy as a parameter of segregation and then 18 phytochemical-protein complexes were created for the purpose of further analysis using Discovery Studio 2019, which was then utilised for identifying the forces of attraction between the phytochemical and the protein, the interacting residues and to produce a 2D diagram of the best complex confirmation for each complex (Figure 3). The entire analysis indicated that although Aphidicolin and Epicatechin (EC) has good results as well; Nuciferine, Retinal and Salvinorin A possess the best results among all the phytochemicals, when considering both SLC6A3 and SLC6A4. On the other hand, while another one of the remaining phytochemicals, Forskolin, had a good score for SLC6A4, it can’t be considered for succeeding works because the purpose of this study is to identify potential drug candidates which can target both SLC6A3 and SLC6A4 at the same time, not just one of them (Table 3).

**Figure 3:**
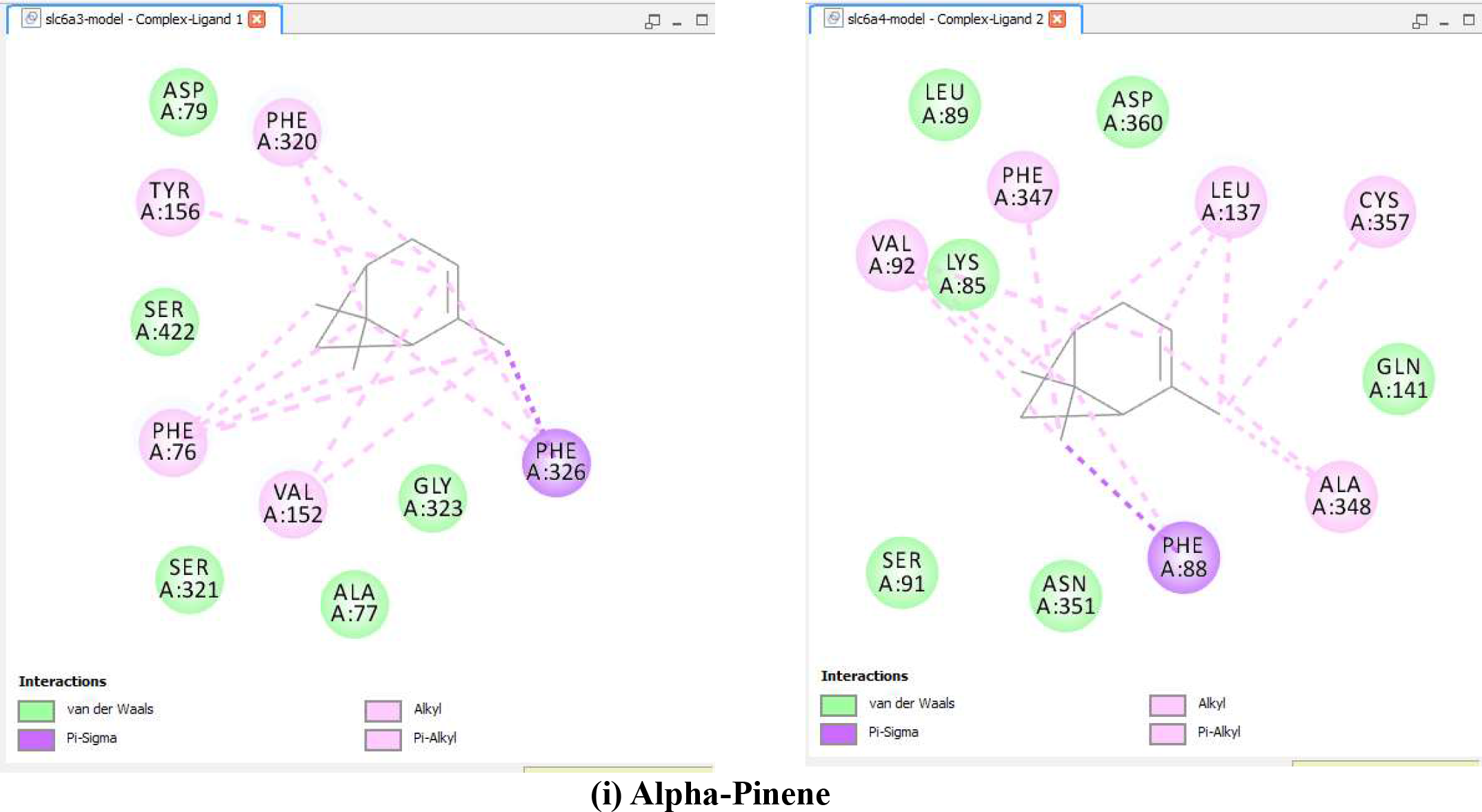
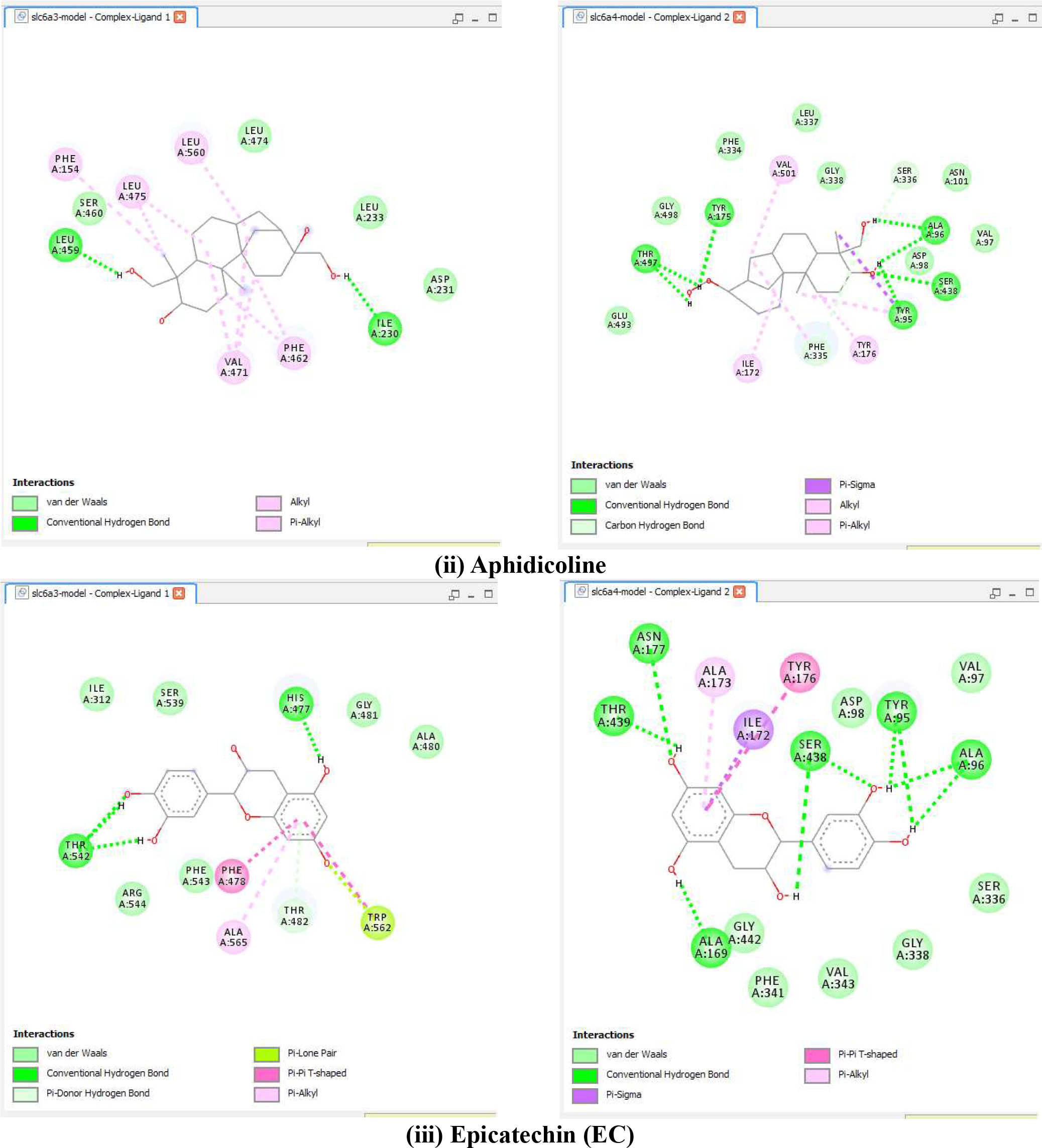
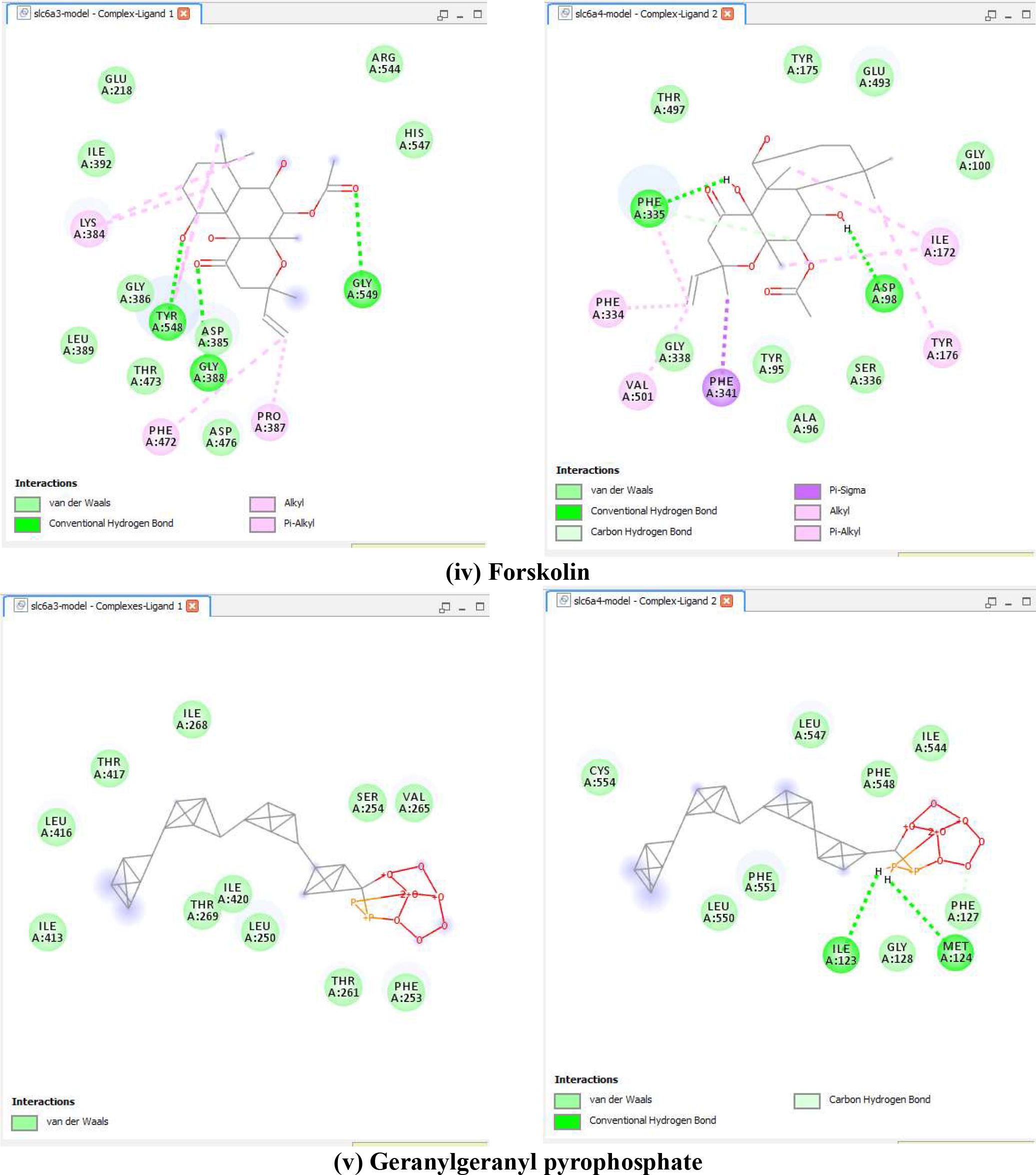
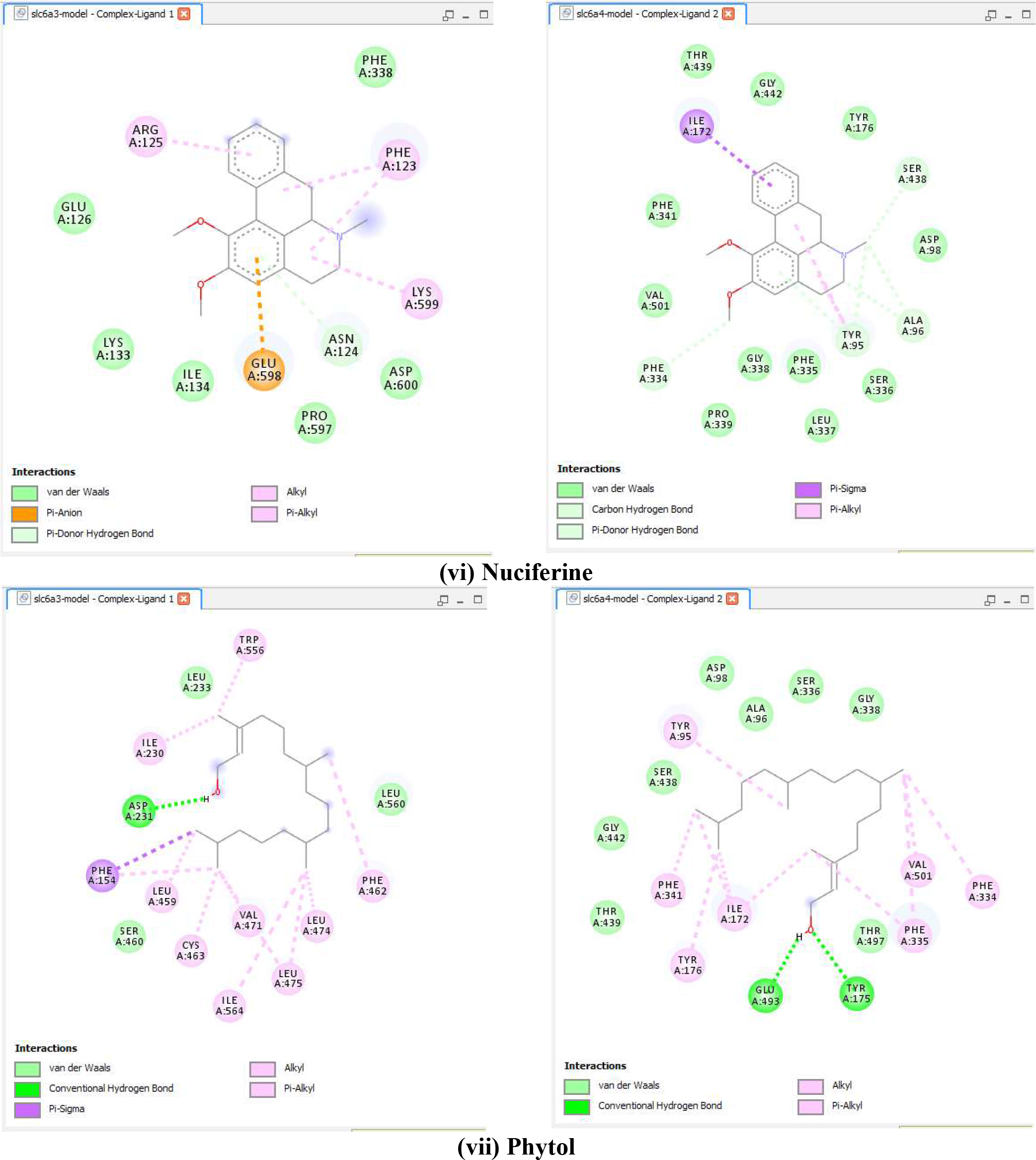
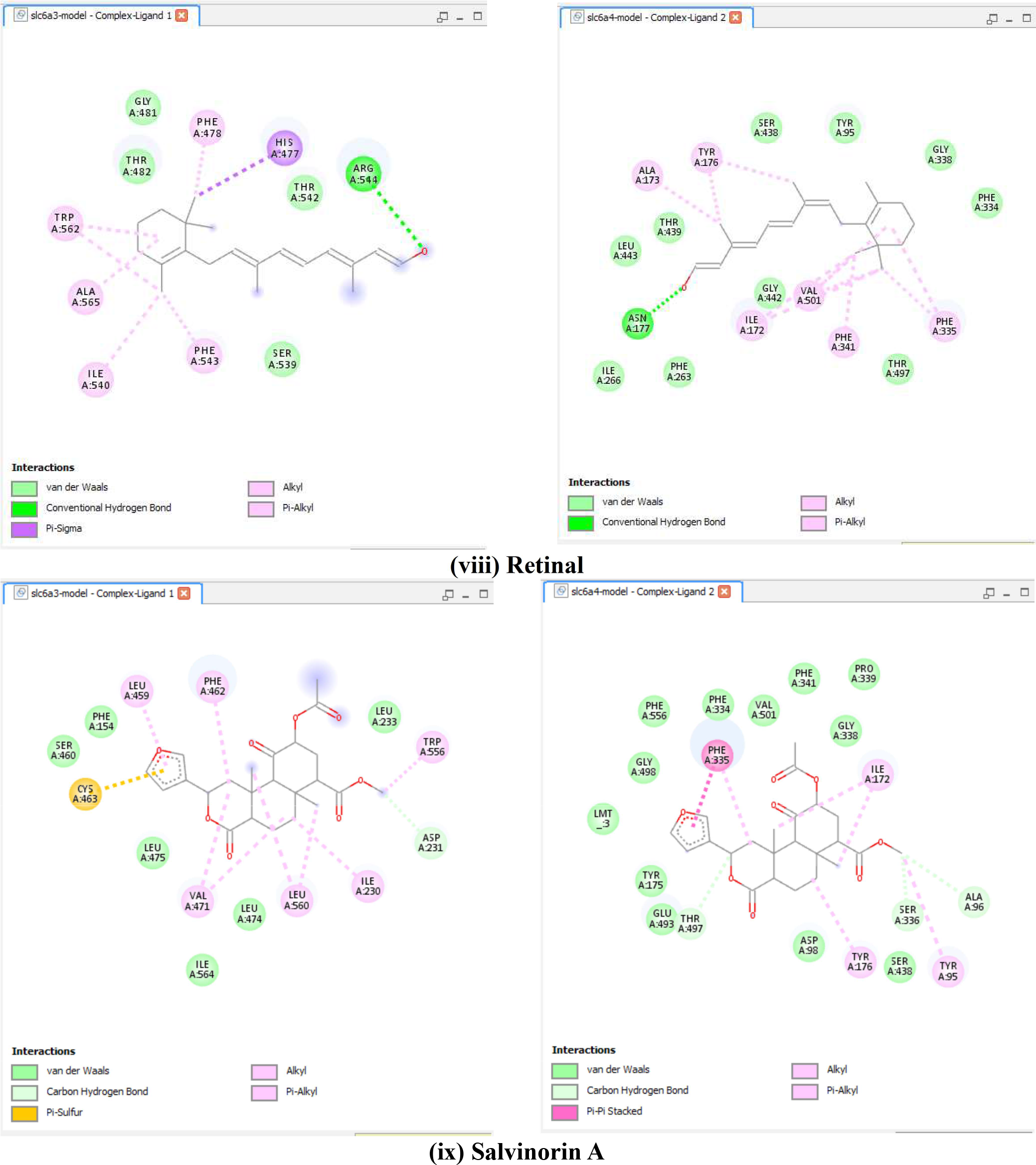
Residue interaction visualisations of best molecular docking configurations for the phytochemical ligands as complexes with SLC6A3 (left) and SLC6A4 (right)

**Table 3:**
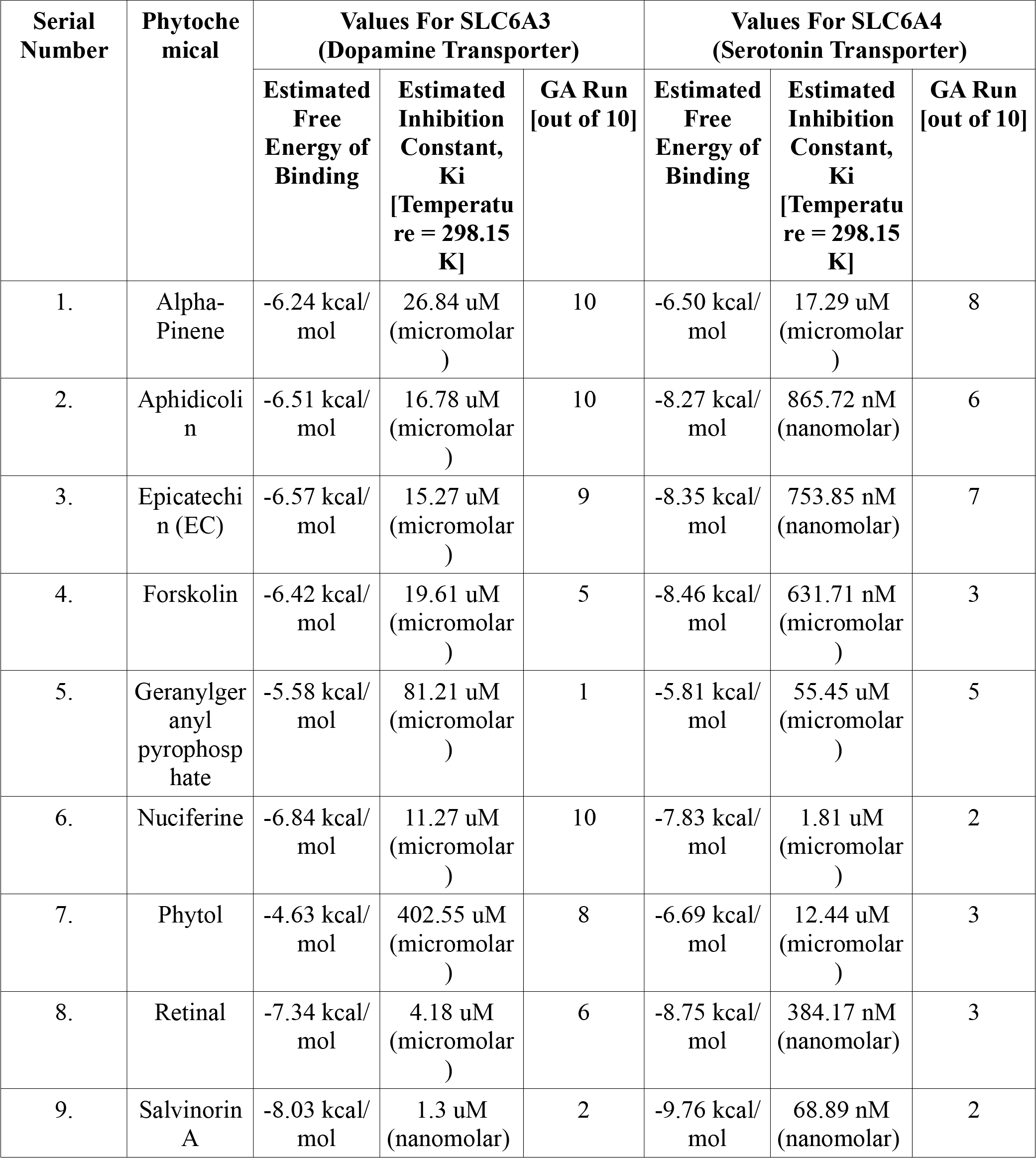
Top molecular docking scores of phytochemical ligands for both SLC6A3 and SLC6A4 out of ten configurations for each.

## CONCLUSION

The current work provides fresh insight, supplementing the previously lacking information about how known phytochemicals can be used as potential drugs for the treatment of OCD through their association with the proteins involved in the expression of its symptoms. While several phytochemicals have shown good molecular docking scores for both the transport proteins, only two phytochemicals: Nuciferin, an alkaloid extract of Indian Lotus plant, which is also known to be present in Blue Egyptian Lotus plant, plus Retinal, also referred to as Retinaldehyde, appear to be the two most suitable compounds on the basis of various factors, such as free binding energy and inhibition constant calculations, for both dopamine transporter and serotonin transporter, along with the fact that Nuciferine is known for inducing sedation; plus inhibiting amphetamine toxicity, conditioned avoidance response, spontaneous motor activity and stereotypy (it could potentiate morphine analgesia as well); while Retinal is known to be a Vitamin A aldehyde [25][26][27][28]. On the other hand, although it shows better molecular docking scores than Nuciferine and Retinal do, Salvinorin A, an extract of the mexican plant Salvia divinorum, appears to be identified as an illegal psychotropic substance in several countries, including Australia and Sweden, due to the considerable number of negative side-effects it shows upon consumption, even in small quantities [29][30]. Hence, we can safely conclude that Nuciferine and Retinal could be good revealing therapeutic agents for OCD and are the best candidates for further research among all the phytochemicals covered in this study.

## ACKNOWLEDGEMENTS

I’d like to acknowledge the creators and maintainers of PubMed, LibGen, PDB, UniProt, Swiss-Model, SOPMA, ProtParam, SAVES, UCSF Chimera, Molinspiration, Wikipedia, PubChem, Open Babel, Open Babel GUI, AutoDock, AutoDockTools and Discovery Studio for the immense help these databases/websites/softwares provided us with during the course of our research.

